# Development of endothelial-targeted CD39 as a therapy for ischaemic stroke

**DOI:** 10.1101/2023.12.12.571382

**Authors:** Natasha Ting Lee, Ioanna Savvidou, Carly Selan, Ilaria Calvello, David K Wright, Robert Brkljaca, Abbey Willcox, Joanne SJ Chia, Xiaowei Wang, Karlheinz Peter, Simon C. Robson, Robert L Medcalf, Harshal H Nandurkar, Maithili Sashindranath

## Abstract

**Background:** Ischaemic stroke is characterized by a necrotic lesion in the brain surrounded by an area of dying cells termed the penumbra. Salvaging the penumbra either with thrombolysis or mechanical retrieval is the cornerstone of stroke management. At-risk neuronal cells release extracellular adenosine triphosphate (eATP) triggering microglial activation and causing a thromboinflammatory response culminating in endothelial activation and vascular disruption. This is further aggravated by ischaemia-reperfusion (I/R) injury that follows all reperfusion therapies. The ecto-enzyme CD39 regulates eATP by hydrolysing to adenosine which has anti-thrombotic and anti-inflammatory properties and reverses I/R injury.

**Methods:** We developed *anti-VCAM-CD39* that targets the antithrombotic and anti-inflammatory properties of recombinant CD39 to the activated endothelium of the penumbra by binding to vascular cell adhesion molecule (VCAM)-1. Mice were subjected to 30 minutes of middle cerebral artery occlusion (MCAo) and analysed at 24h. *Anti-VCAM-CD39* or control agents (saline, non-targeted CD39, or anti-VCAM-inactive CD39) were given at 3h post-MCAo.

**Results:** Anti-VCAM-CD39 treatment reduced neurological deficit; MRI confirmed significantly smaller infarcts together with an increase in cerebrovascular perfusion. Anti-VCAM-CD39 also restored blood brain barrier (BBB) integrity and reduced microglial activation. Coadministration of *anti-VCAM-CD39* with thrombolytics (tPA) further reduced infarct volumes and attenuated BBB permeability with no associated increase in intracranial haemorrhage.

**Conclusion:** *Anti-VCAM-CD39*, uniquely targeted to endothelial cells, could be a new stroke therapy even when administered 3 h post ischaemia and may further synergise with thrombolytic therapy to improve stroke outcomes.

## Introduction

Ischaemic stroke, which results from transient or permanent reduction in cerebral blood flow (cBF) of a major brain artery(1, 2), is caused by thrombosis or thromboembolism. A debilitating neurological condition, it is the second leading cause of death worldwide(3, 4) and is one of the major causes of disability in Australia(5). Injury to the brain is multi-faceted – during ischaemia, cessation of blood flow causes oxygen-glucose deprivation leading to metabolic failure and cell death(6). The reestablishment of blood flow after clot dissolution or retrieval paradoxically leads to ischaemic reperfusion (I/R) injury(7) triggering the release of reactive oxygen species (ROS) that activates microglia and astrocytes, as well as increases the expression and release of pro-inflammatory cytokines TNFα, IL-1β and IL-18(3).

Inflammation and oxidative stress trigger endothelial dysfunction. Endothelial cells are a critical structural component of the neurovascular unit that constitutes the blood-brain barrier (BBB). Endothelium dysfunction and death after I/R injury not only promotes neuroinflammation and brain oedema, but also increases the risk of intracerebral haemorrhage of thrombolytic therapies(8, 9). Although timely recanalization of the occluded vessel can aid in salvaging the ischaemic zone, the ischaemic penumbra is the area where pharmacologic interventions are most likely to be effective(10). This zone is comprised of activated endothelial cells and dying neurons; it has preserved energy metabolism and is salvageable for ∼6-8 hours after stroke(11). Within the first 24 hours, there is a broader area around the initial ischaemic core and wider than the ischaemic penumbra. Known as the inflammatory penumbra, it is characterised by abundant endothelial activation with VCAM-1 expression(12). In response to ischaemic injury, ICAM-1 and VCAM-1 levels on brain endothelial cells are upregulated 3-15-fold(13–16), reaching maximum levels at 12 to 24 hours post injury. Immune cells, especially T-cells and monocytes, bind to endothelial VCAM-1 which allows them to migrate across the endothelium into the brain. This further aggravates cell death and eventually the inflammatory penumbra also becomes part of the necrotic core(12).

Following focal ischaemia, ATP is rapidly released into the extracellular space by dying neurons and persists for up to 24 hours(17). The source of extracellular ATP (eATP) is dying neurons within the infarcted tissue that in turn activate microglia, most likely via P2X7 receptors(18) triggering microglial chemotaxis. Microglial P2X7 receptor activation leads to the nuclear translocation of NF-κB (nuclear factor kappa-light-chain enhancer of activated B cells), that increases expression of TNF-α and IL1β(19). P2X7 activation is also associated with formation of the NLRP3 inflammasome in macrophages and microglia, characterized by IL1β/IL-18 release and activation of caspase-1 that promotes neuronal apoptosis(18). In stroke, activated microglia accumulate in the vicinity of the infarct, correlating with an increase in TNF-α in the brain within 1 to 3 hours of injury(20, 21). Microglial TNF-α activates endothelial cells and can promote endothelial necroptosis after MCAo(22).

While eATP is pro-inflammatory, ADP activates platelets and promotes thrombosis. eATP and ADP are hydrolysed to AMP by CD39 that is anchored on the endothelial cell surface. AMP is subsequently hydrolysed to adenosine by CD73, another endothelial cell surface enzyme. Adenosine has opposing actions to ATP/ADP: adenosine is anti-inflammatory in contrast to the pro-coagulant/pro-inflammatory actions of ADP and ATP. There is a loss of CD39 activity during endothelial dysfunction(23) and therapeutic delivery of CD39 is protective in ischaemic stroke(24). However, a prevailing issue with systemic CD39 therapeutics is disruption of haemostasis, because of impaired systemic platelet aggregation which promotes bleeding(25–27). In contrast, targeted-CD39 therapies promote localized, sustained ATP/ADP hydrolysis and adenosine generation at the site of cellular injury without high systemic concentrations associated with bleeding effects (28, 29).

Here we explored the utility of a novel therapeutic, *anti-VCAM-CD39* which consists of recombinant CD39 fused to an anti-VCAM-1 single-chain variable fragment fusion protein (scFv). We hypothesise that this bifunctional construct would confer the anti-inflammatory and antithrombotic effect of CD39-catalysed adenosine generation to areas of endothelial dysfunction characterised by upregulation of VCAM-1.

## Results

### *Anti-VCAM-CD39* was shown to be fully functional and decreased overall *in vitro* leukocyte transmigration across an activated endothelial cell monolayer

To evaluate the effect of the VCAM-1-ScFv component of *anti-VCAM-CD39* we evaluated the migration of bone marrow-derived immune cells through an endothelial monolayer after exposing the cells to IL-1β and TNFα, which are pro-inflammatory cytokines that are commonly upregulated by endothelial cells after damage. We found that overall cell migration was decreased after the endothelial layer was treated with *anti-VCAM-CD39* following 4 hours of IL-1β pretreatment (not shown). Identification of the individual migrated cell populations was done by trained pathologist blinded to the groups, and we found that the decrease was most pronounced in the myeloid cell population, where pretreatment with IL-1β significantly increased cell migration and following administration of *anti-VCAM-CD39* transendothelial migration was significantly limited. The protection observed by the non-targeted CD39 construct suggests that soluble CD39 could itself be protecting the endothelial monolayer by promoting adenosine generation (**Supplementary Figure A**).

Overall cell migration was significantly decreased when the endothelial layer was treated with *anti-VCAM-CD39* after 24 hours of TNFα pretreatment, specifically in the lymphocyte and myeloid populations (**Supplementary Figure B**). Further, we found that the non-targeted VCAM-CD39 was not able to reduce cell migration confirming that VCAM blockade rather than CD39 activity was involved in reducing TNFα induced inflammatory cell migration.

We also found that tail bleeding was only significantly increased at a dose of 2mg/kg (**Supplementary Figure C**). Further, administration of non-targeted CD39 at 1-1.5mg/kg did not increase tail bleeding time. Taken together, this suggests that *anti-VCAM-CD39* is safe to administer at concentrations below 2mg/kg.

### *Anti-VCAM-CD39* ameliorates brain damage in mice subjected to MCAo

MRI was performed 24hours after mice underwent MCAo and treatment with 0.5mg/kg of *anti-VCAM-CD39,* control agents or saline. We found that there was a ∼40% decrease in infarct size, demonstrating the neuroprotective effect of *anti-VCAM-CD39*. (Saline: 85.22±7.828 mm^3^, n=7; *anti-VCAM-CD39*: 51.08±5.3 mm^3^, n=6, p=0.0137) (**Figure 1A**). The protection by *anti-VCAM-CD39* was superior to the effects of non-targeted and inactive CD39 controls, which despite causing some reduction in infarct volume compared to saline treated animals yielded a large variation in infarct volumes. This partial protection is expected, due to the independent activities of the anti-VCAM ScFv component blocking transendothelial immune cell migration and systemic ATP/ADP hydrolytic effects of non-targeted CD39.

**Figure 1:**
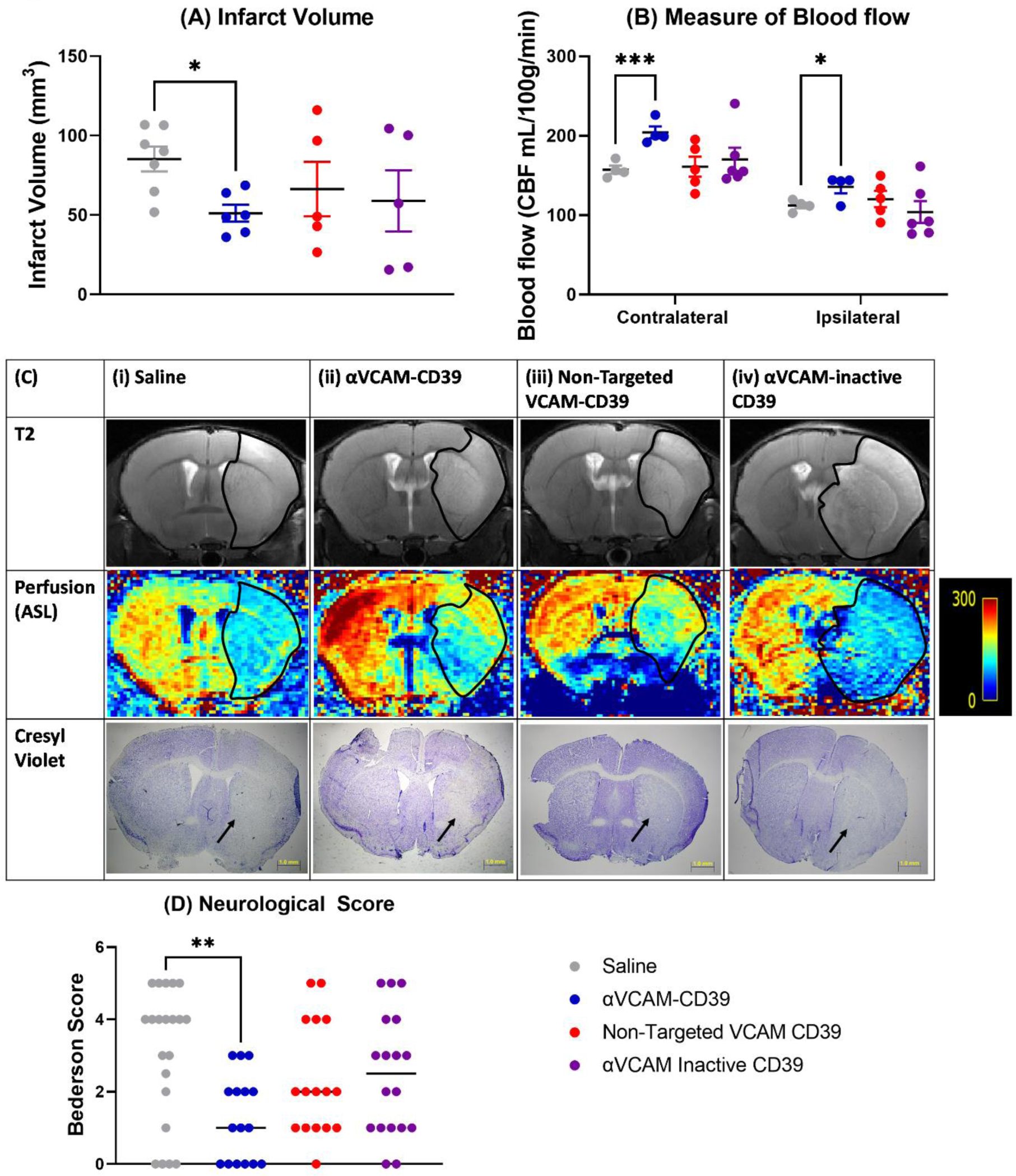
*Anti-VCAM-CD39* is effective in protecting the brain in mice post-MCAo. **(A)** MRI-based DWI-quantified infarct volumes (mm^3^) of *anti-VCAM-CD39* treated mouse brains 24 hours post-MCAo showed significantly lower infarct volumes compared to saline, non-targeted VCAM-CD39 and *anti-VCAM-inactive CD39* controls. (Data is Mean ± SEM; Saline: n=7, Treated: n=5-6; *p<0.05; Brown-Forsythe and Welch ANOVA test with Dunnett’s T3 post-hoc analysis). **(B)** Cerebrovascular perfusion (presented as intensity) was increased in both hemispheres after *anti-VCAM-CD39* treatment (Data is Mean ± SEM; Saline: n=4, Treated: n=4; *p<0.05, ***p<0.001; two-way ANOVA with Sidak’s post-hoc analysis). **(C)** MR T2 imaging, MR perfusion imaging and Cresyl Violet staining of mouse brains post-MCAo after treatment with (i) saline, (ii) *anti-VCAM-CD39*, (iii) non-targeted VCAM-CD39, (iv) *anti-VCAM-inactive CD39* (n=4-5) distinctly show the infarcted area indicated by delineation in the T2 imaging, and the corresponding perfusion image showing increased amount of blood flow within and around the infarct after *anti-VCAM-CD39* treatment, perfusion mean denoted by the colour scale on the right. This injury and corresponding absence of Cresyl violet positive cells in the same region, indicated by the arrows, confirms cell death within the infarct. **(D)** *Anti-VCAM-CD39* significantly reduced neurological deficit compared to saline, and also had lower scores compared to control groups (Sham: n=12, Saline: n=21, Treated: n=16-18; Bars indicate median value, **p<0.01, ****p<0.0001; One-way ANOVA test with Dunnett’s post-hoc analysis).

Arterial spin labelling (ASL - tissue perfusion) was utilised as a measure of relative CBF (measured in mL/100g of tissue/minute) in selected areas of the brain, predicted to span the core and therefore constitute penumbra(30). We found that with the administration of *anti-VCAM-CD39*, there was a significant increase in CBF in both the left and right hemisphere (-2mm from bregma) (**Figure 1B**). However, unlike *anti-VCAM-CD39,* non-targeted VCAM-CD39 and *anti-VCAM-inactive CD39* did not significantly increase blood flow on either side of the brain.

Representative MRI and Cresyl violet histology images of mice post-MCAo support the findings that with *anti-VCAM-CD39* treatment, infarct size was significantly decreased (**Figure 1C**). There was overall increased blood flow within the brain after *anti-VCAM-CD39* treatment, corresponding to an increase in the perfusion mean when compared to saline treated mice. The absence of Cresyl violet positive cells in the infarcted area in corresponding tissue sections confirmed these findings. Treatment with non-targeted VCAM-CD39 and *anti-VCAM-inactive CD39* was less effective in reducing infarct volume, and increasing blood flow, when compared to *anti-VCAM-CD39*, evidenced by a larger delineated infarct, and lower cBF, denoted by a lower intensity perfusion signal within the perfusion maps. As the MCAo model does not involve an occlusive thrombus and is based on I/R injury, this suggests that the reduction in infarct volume and increase in cBF by *anti-VCAM-CD39* is conferred through the bifunctional action of targeting VCAM-1 and therapeutic benefit of CD39 catalytic activity localised to the microcirculation.

Behavioural function assessed by an individual blinded to the groups at 24 hours post-MCAo showed a significant increase in neurological deficit in mice post-MCAo (p=0.0001) that was abrogated after *anti-VCAM-CD39* treatment (**Figure 1D**). Administration of non-targeted VCAM-CD39 and *anti-VCAM-inactive CD39* treatment did not substantively alter neurological deficit.

### *Anti-VCAM-CD39* suppresses inflammation, cell death and increases adenosine generation in mice post-MCAo

We found that *anti-VCAM-CD39* treatment significantly downregulated gene expression of several inflammatory cytokines, including IL-1β (**Figure 2A**), IL-18 (**Figure 2B**) and TNFα (**Figure 2C**), suggesting an overall reduction of inflammation post-treatment. Additionally, we found significant decrease in CXCL1 expression relative to saline treated mice (**Figure 2D**), which could indicate a reduction in neutrophil and monocyte infiltration(31).

**Figure 2:**
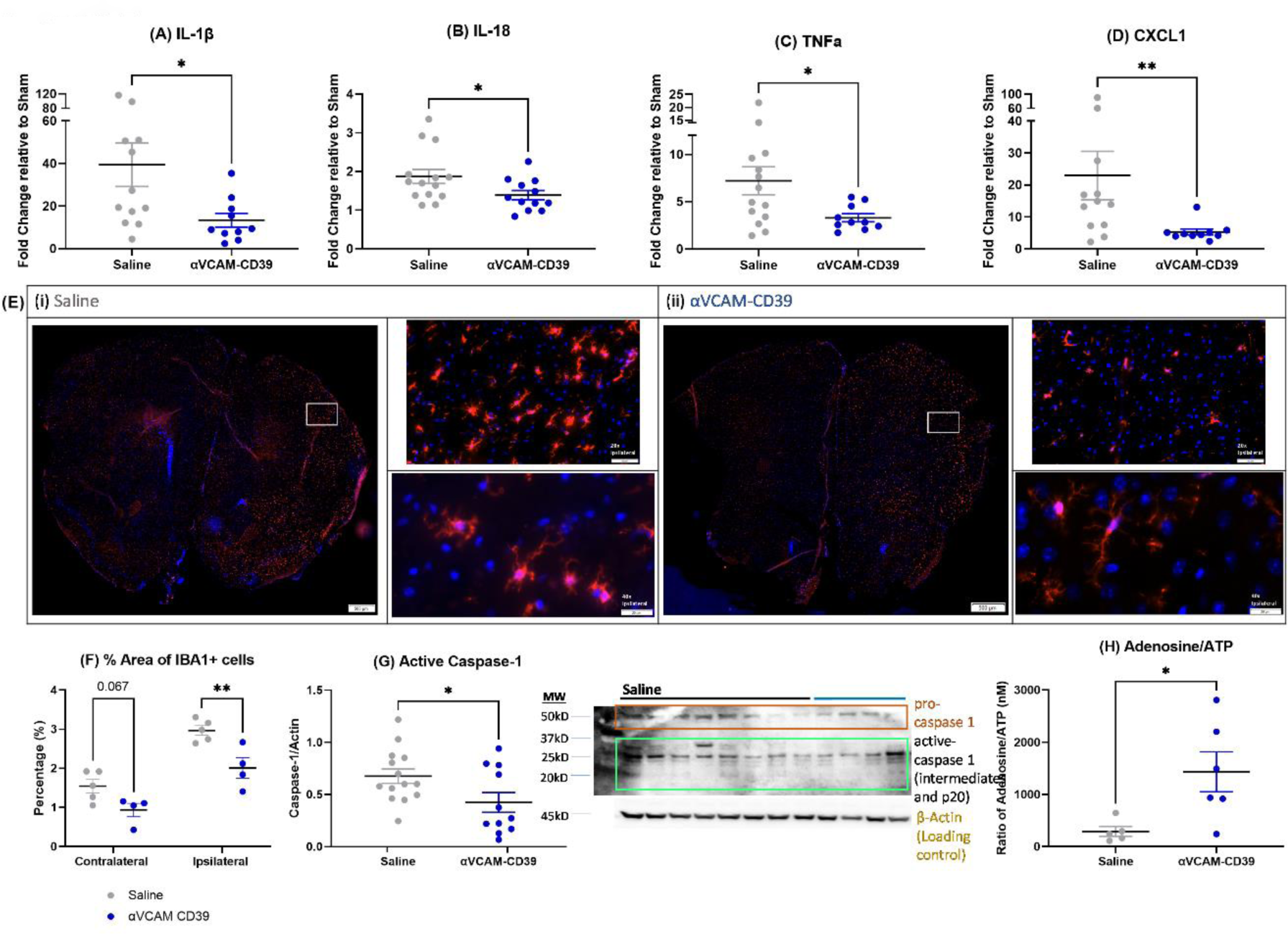
Gene expression analysis of chemokines and inflammatory cytokines, immunohistochemistry of microglia, protein expression of cleaved caspase-1 and ratio of systemic concentration of Adenosine/ATP showed significant changes after administration of *anti-VCAM-CD39* post-MCAo. Real time PCR analysis of RNA isolated from the site of injury at 24 hours shows significant changes in the gene expression. Expression of inflammatory cytokines **(A)** IL-1β, **(B)** IL-18 and **(C)** TNFα were significantly decreased after *anti-VCAM-CD39* treatment. Chemokines **(D)** CXCL1 was significantly decreased in expression after *anti-VCAM-CD39* treatment. (Data is Mean ±SEM; Saline: n= 12-14, 0.5mg/kg *anti-VCAM-CD39*: n=10-12; *p<0.05; Unpaired t-test). **(E)** Percentage of area with IBA1 positive cells are significantly reduced after *anti-VCAM-CD39* treatment post-MCAo. (Data is Mean ± SEM; Saline: n=5, Treated: n=4; **p<0.01; two-way ANOVA with Sidak’s post-hoc analysis). **(F)** IBA1 is visibly significantly decreased in the (ii) treated cohort compared to the (i) saline treated group. Magnification at 20x at the ipsilateral side shows that after *anti-VCAM-CD39* treatment, the microglia appear ramified, while microglia in the saline treated group appear activated, and amoeboid in shape. 40x magnification show the distinctly different microglial morphology post *Anti-VCAM-CD39* treatment. indicative of reduced diseased state (Immunostaining with IBA1 to delineate microglia (red) and DAPI to identify nuclei. Representative images of N=8 per mouse, and n=4-5 per group. **(G)** Caspase-1 activity was found to be significantly decreased after *anti-VCAM-CD39* treatment post-MCAo (Data is Mean ±SEM; Saline: n= 14, *anti-VCAM-CD39*: n=11; *p<0.05; Unpaired t-test). **(H)** Systemic Adenosine and ATP concentration (expressed as a ratio) in plasma was found to be significantly increased after *anti-VCAM-CD39* treatment (Data is Mean ±SEM; Saline: n=5, *anti-VCAM-CD39*: n=6; *p<0.05; Unpaired t-test).

Microglial activation was significantly reduced after *anti-VCAM-CD39* treatment (**Figure 2E**), evidenced by an overall decrease in the area occupied by IBA1+ cells relative to the area of each hemisphere (p=0.0048) mirroring a reduction in size of these microglia (**Supplementary Figure D**). These data validate the visual observations (**Figure 2F**), that the microglia had a more quiescent, ramified morphology(32) in the brains of anti-VCAM-CD39 treated animals.

Further, *anti-VCAM-CD39* treatment significantly reduced the expression of cleaved caspase-1 (**Figure 2G**) and reduced gene expression of IL-1β and IL-18. The shift in the plasma adenosine/ATP ratio confirms the *in vivo* enzymatic activity of *anti-VCAM-CD39* to reduce eATP in the systemic circulation and leads us to postulate that a more profound effect would occur in the targeted cerebral ischaemic microcirculation. (**Figure 2H**). Together, these data strongly indicate that *anti-VCAM-CD39* may protect the brain by reducing eATP and consequent activation of the NF-κB/NLRP3 inflammasome pathway(33–36).

### *Anti-VCAM-CD39* reduced blood brain barrier permeability and limited endothelial activation

We confirmed a significant increase in albumin extravasation in saline treated mice post-MCAo (**Figure 3A**) suggesting significant BBB breakdown. BBB integrity was restored with *anti-VCAM-CD39* treatment with no significant improvement noted for the administration of non-targeted and inactive CD39 controls. Release of vWF is a feature of endothelial activation(37) and consistent with induction of I/R injury, plasma vWF increased post-MCAo when compared to sham (as seen in the representative western blot). Treatment with *anti-VCAM-CD39* (**Figure 3B**) significantly reduced plasma vWF, thereby confirming its endothelial protective actions.

**Figure 3:**
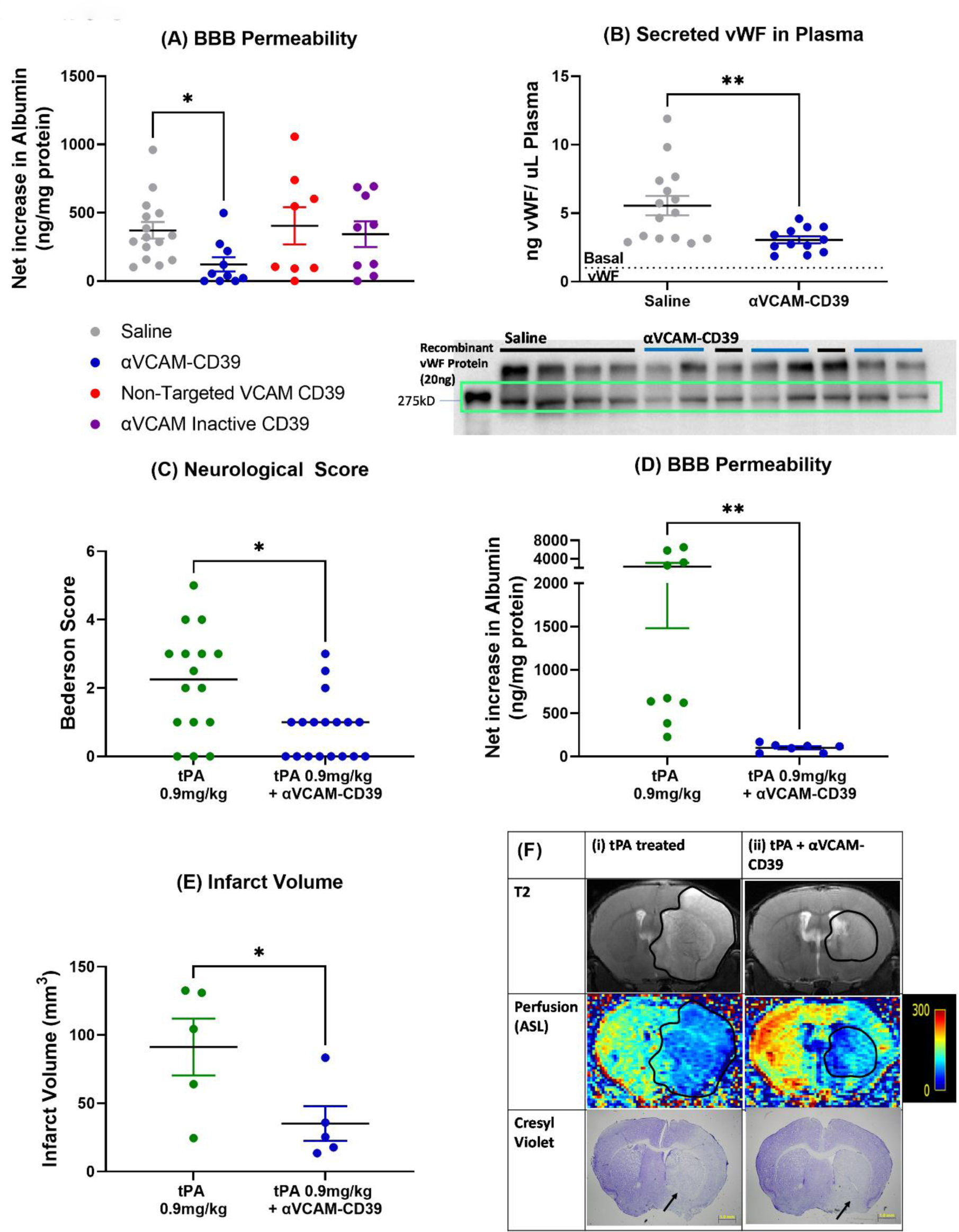
*Anti-VCAM-CD39* significantly decreased blood brain barrier permeability and systemic vWF levels post-MCAo, and was effective in protecting the brain after tPA treatment in mice post-MCAo. **(A)** BBB permeability measured by net albumin extravasation in the brain parenchyma was significantly increased post-MCAo, and *anti-VCAM-CD39* treatment was effective in reducing albumin extravasation (Data is Mean ±SEM; Saline: n=15, Treated: n=8-10; *p<0.05; Kruskal-Wallis test, Dunn’s multiple comparisons test). **(B)** Circulating vWF in plasma was found to be significantly reduced after *anti-VCAM-CD39* treatment (Data is Mean ±SEM; Saline: n=15, Treated: n=12; **p<0.01; unpaired t-test). **(C)** *Anti-VCAM-CD39* after tPA treatment significantly reduced neurological deficit compared to the tPA treated group (tPA alone: n=16, *Anti-VCAM-CD39*: n=18; Bars indicate median value, *p<0.05; One-way ANOVA with Uncorrected Fishers LSD). **(D)** *anti-VCAM-CD39* treatment was effective in reducing net BBB permeability was measured by albumin extravasation (Data is Mean±SEM; tPA: n= 9, tPA + Anti-VCAM-CD39: n=7; Kruskal-Wallis test, Dunn’s multiple comparisons test). **(E)** MRI-based DWI-quantified infarct volumes (mm^3^) of tPA + *anti-VCAM-CD39* treated mouse brains 24 hours post-MCAo showed significantly lower infarct volumes compared to tPA treated, (Data is Mean ± SEM; tPA: n=5, *Anti-VCAM-CD39*: n=5; *p<0.05; One-way ANOVA with Uncorrected Fishers LSD). **(F)** MR T2 imaging, MR perfusion imaging and Cresyl Violet staining of mouse brains post-MCAo after treatment with (i) tPA, and (ii) *anti-VCAM-CD39*, (n=5) distinctly show the infarcted area indicated by delineation in the T2 imaging, and the corresponding perfusion image showing increased amount of blood flow within and around the infarct after *anti-VCAM-CD39* treatment. This injury and corresponding absence of Cresyl violet positive cells in the same region, indicated by the arrows, confirms cell death within the infarct.

### *Anti-VCAM-CD39* synergised with tPA in improving outcome after MCAo

Given that the MCAo model reflects I/R injury rather than ischaemia after acute embolic stroke(38, 39), tPA treatment alone was able to modestly reduce neurological deficit (p=0.07) reflecting its possible effect on microcirculatory thrombosis due to endothelial activation (**Figure 1D and 3C**). Co-administration of *anti-VCAM-CD39* with tPA significantly decreased functional neurological deficit when compared to saline treated mice as well as mice with tPA treatment alone (**Figure 3C**). Albumin extravasation in the ischaemic ipsilateral side of the brain was increased four-fold after tPA treatment. However, co-administration of *anti-VCAM-CD39* with tPA, led to a significant decrease in albumin extravasation confirming its protective effect on BBB disruption even in the presence of tPA (**Figure 3D**).

tPA treatment resulted in infarcts of similar volume to saline treated mice but co-administration of *anti-VCAM-CD39* led to a significant decrease in infarct size compared to saline and tPA treated mice. (**Figure 3E**). MRI showed that tPA treatment alone after MCAo resulted in widespread infarcts throughout the hemisphere similar to saline treated mice, but with tPA + *anti-VCAM-CD39* treatment, there was a significant decrease both in infarct size and in midline distortion, suggesting reduced oedema(40, 41) (**Figure 3F**). Corresponding Cresyl Violet stained sections taken from the same brains show a smaller area of cell loss after tPA + *anti-VCAM-CD39* treatment (**Figure 3F**).

### *Anti-VCAM-CD39* did not increase bleeding in an intracerebral haemorrhage model

To provide preclinical data for the safety of *anti-VCAM-CD39* under haemorrhagic stroke conditions i.e. without the need for prior neuroimaging, we used a model of intracerebral haemorrhage (ICH). Systemic administration of *anti-VCAM-CD39* (0.5mg/Kg) after unilateral intracerebral collagenase injection did not worsen neurological outcome (**Figure 4A**) or promote intracerebral bleeding (**Figure 4B**). Interestingly, *anti-VCAM-CD39* was able to reduce apoptotic activity by 30% in the ipsilateral hemisphere with intracerebral collagenase-mediated injury (**Figure 4C**; p=0.005).

**Figure 4:**
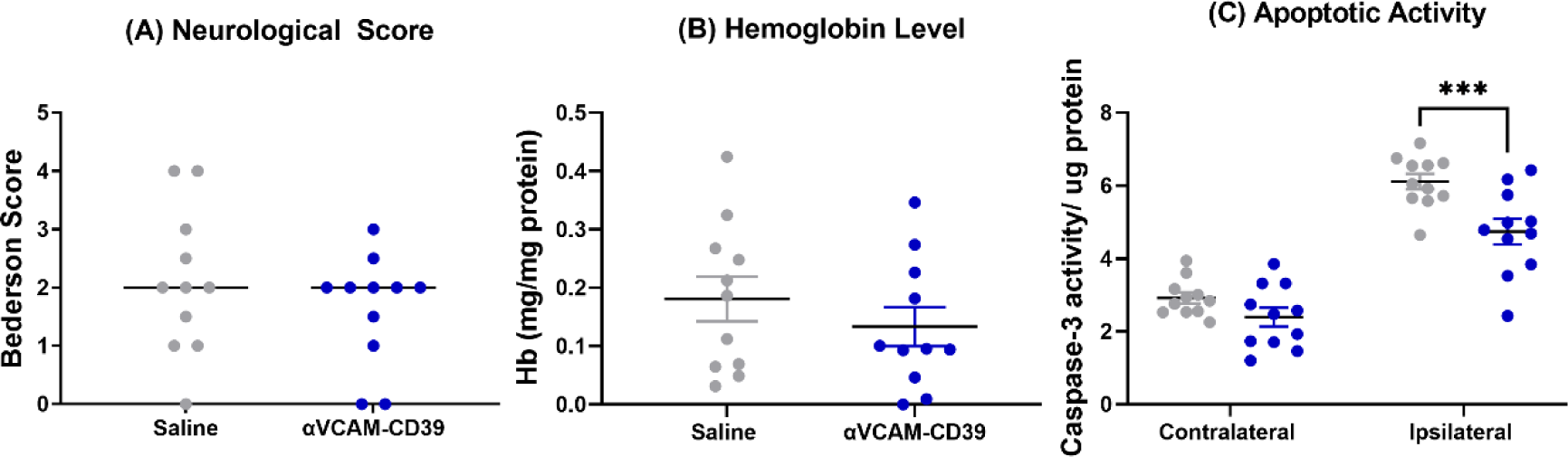
*Anti-VCAM-CD39* did not exacerbate bleeding or worsen brain damage in an intracerebral haemorrhage (ICH) model. *Anti-VCAM-CD39* after ICH did not worsen **(A)** neurological deficit or **(B)** promote bleeding, measured by hemoglobin quantification (Data is Mean±SEM; Saline: n=11, *anti-VCAM-CD39*: n=11; Unpaired t-test). **(C)** Apoptotic activity 24-hours post-ICH after *anti-VCAM-CD39* was shown to be decreased on the ipsilateral side of the brain (Data is Mean±SEM; Saline: n=11, *anti-VCAM-CD39*: n=11; ***p<0.001; Two-Way ANOVA with Uncorrected Fishers LSD).

## Discussion

Here, we investigated the ability of a novel bifunctional construct - *anti-VCAM-CD39* to reduce brain damage and neurological deficits in mice after focal ischaemia.

Endothelial VCAM-1 expression after ischaemia has been well characterised by us and others in both human and rodent models of ischaemic stroke(12, 42–45), and this was the basis of our endothelial targeting strategy for CD39 delivery to the brain. Previous literature has shown that exogenous addition of CD39 at 3 hours post stroke was able to reduce thrombosis, increase post-ischaemic perfusion and decrease cerebral infarct volumes(24). We have also previously characterised the temporal profile of VCAM-1 upregulation in the brain after MCAo(42) and found that the highest upregulation is at 3 hours. Together with the average time required for presentation to hospital and the therapeutic time window for thrombolysis(46), we administered *anti-VCAM-CD39* at 3 hours post-stroke.

Other than being an effective targeting strategy, our *in vitro* data suggests that the blocking capacity of the ScFv VCAM-1 component could limit transendothelial migration of inflammatory cells, specifically microglia and monocytes. However, blocking VCAM-1 alone is not considered sufficient to protect the brain after stroke(47) and our data showing a lack of cerebroprotection with the anti-VCAM-inactive CD39 confirms these findings.

Baek et al showed that mice with transgenic overexpression of human CD39 (hCD39), both globally and restricted to myeloid lineages had smaller infarct volumes after MCAo when compared with wild-type mice and interestingly, CD39 transgenic mice had less macrophage and neutrophil infiltration(48). We demonstrated that *anti-VCAM-CD39* decreased CXCL1 expression; CXCL1 expression by monocytes and neutrophils is known to be increased after ischaemia(31), promoting leukocyte infiltration. We confirmed the enzymatic activity of anti-VCAM-CD39 using ATP degradation and malachite green assays (not shown) and by showing that anti-VCAM-CD39 promoted tail bleeding at doses >2mg/kg. Taken together, it is plausible that the combination of both VCAM blockade and local CD39 delivery is likely to significantly restrict inflammatory cell infiltration after MCAo(48), although this remains to be verified.

Targeting CD39 affords cerebroprotection at a lower dose of 0.5 mg/kg, which does not alter haemostasis. Pinsky et al earlier showed that soluble untargeted CD39 administered pre-operatively inhibited platelet and fibrin accumulation post-MCAo(24), but at significantly higher doses (4-8 mg/kg). These data also confirm the generation of microvascular thrombi in the MCAo model where stroke is induced by ligation rather than thrombosis within the MCA(24). Pinsky et al(24) showed that untargeted solCD39 improved post-ischaemic cerebral perfusion indicating that CD39 can mitigate microvascular thrombosis by reducing ADP-mediated platelet activation as well as endothelial activation(24). Additionally, increased cBF correlated with reduced infarct volumes(24) which is akin to our findings showing improved cerebrovascular perfusion and lower infarct volumes in *anti-VCAM-CD39* treated animals. The non-targeted CD39 construct that we used functions like solCD39; unsurprisingly, we saw some protection with this construct but there was a high degree of variability owing to the dose being lower than that used by Pinsky et al(24). The MCAo model does not have an occlusive thrombus so the only detected increase in CBF was due to improved microvascular perfusion and in fact, this is highly relevant to rescuing the ischaemic penumbra. Therefore these results further validate that targeting CD39 to the ischaemic penumbra characterised by microthrombosis and endothelial activation is an important therapeutic strategy for ischaemic stroke.

We used RT-PCR to quantify cytokine changes within the ischaemic tissue as we are interested in detecting changes in neuroinflammation following anti-VCAM-CD39 treatment rather than systemic cytokine profiles typically detected in plasma samples. Expression of TNF-α, IL1β, IL-18 was reduced, concomitant with diminished caspase 1 protein levels and reduced microglial activation in the ipsilateral hemispheres of brains of mice subjected to MCAo and treated with *anti-VCAM-CD39*. These findings support the notion that eATP catabolism would reduce P2X7 receptor signaling(49, 50), microglial activation(47) and NLRP3 formation, thereby reducing neuroinflammation(51, 52). However, in animal studies, P2X7 inhibition or gene deletion alone does not consistently promote neuroprotection after MCAo(53). Although P2X7 inhibition has not yet been trialled for cardiovascular conditions, blocking P2X7-IL1β and IL-18 is not effective in suppressing inflammation in a rheumatoid arthritis clinical trial, where pathological P2X7 activation is implicated(54). The advantage of *anti-VCAM-CD39* is that in addition to ATP hydrolysis, the observed increase in cBF suggests that it also sustains adenosine generation. While we have not investigated which adenosine receptor is involved in the protective functions seen in our study, it is known that A_2B_ receptor (A_2B_R) signalling is important for controlling endothelial barrier function(55). During hypoxia, A_2B_R knockout mice have excess vascular leakage in several organs including the brain and expressed higher levels E-selectin and P-selectin in the peripheral vasculature(55). Under basal conditions, A_2B_R deficient mice showed increased adherence of leukocytes to the endothelial layer and increased transendothelial migration of these cells in mesenteric arteries(56). These data support our hypothesis that the cerebroprotective properties of *anti-VCAM-CD39* that we have observed are partially due to increased adenosine generation promoting A_2B_R activation and downstream signalling, although further experiments are needed to confirm this.

Endothelial cells form the luminal compartment of the BBB and therefore, when the endothelium is disrupted, the BBB is compromised. *Anti-VCAM-CD39* significantly reduced BBB damage, likely due to its mitigative actions on platelet activation and thromboinflammation contributing to endothelial protection. We have shown that eATP levels are reduced with anti-VCAM-CD39. The reduction in secreted vWF shows that endothelial activation that aggravates the pro-thrombotic response is reduced with anti-VCAM-CD39, consistent with the dual hydrolytic effects of CD39 on both ATP and ADP.

tPA, commercially known as Alteplase, is a serine protease that activates plasminogen into plasmin, a fibrinolytic enzyme that breaks down fibrin-based blood clots(57). While thrombolysis is the gold standard of care for ischaemic stroke, endovascular thrombectomy is also increasingly used where the infrastructure is available. Our intention was not to validate the synergy between tPA and anti-VCAM-CD39 in their thrombolytic capacity. In both instances, eligibility criteria are strict, and for eligible patients, despite successful recanalization, the infarcted area often increases in size due to I/R injury(58–60). With tPA thrombolysis, approximately 30% of patients have bleeding in the brain (haemorrhagic transformation) which is strongly linked with BBB disruption(61). We are interested in determining whether anti-VCAM-CD39 will reduce the known effects of tPA on intracerebral haemorrhage by attenuating ischemia-reperfusion injury (IRI). Our model characterised the effects of filament-induced acute ischaemia followed by reperfusion injury. This model is widely used in preclinical stroke research and is a true model of ischemia-reperfusion injury (IRI) and there was no occlusive thrombus. It is well known that unlike the MCAo model, endothelial damage is minimal in the thromboembolic models, which would make these models unsuitable for addressing our hypothesis.

After MCAo, the lesion volume was comparable between saline and tPA treated mice, consistent with the literature; since MCAo is an I/R injury model rather than a thrombotic model that is amenable to thrombolysis(38, 39). However, co-administration of anti-VCAM-CD39 with tPA caused a significant improvement in lesion volume and neurological score, at a greater degree than anti-VCAM-CD39 alone (anti-VCAM-CD39: 51.08±5.3 mm3; tPA+ Anti-VCAM-CD39: 35.24±12.65 mm3). This suggests that the restoration of peri-infarct microvascular perfusion with anti-VCAM-CD39 could be enhanced with tPA co-administration while the endothelial-protective actions of CD39 could simultaneously restrict BBB damage associated with tPA treatment. In our stroke model we were unable to detect any haemorrhagic transformation when we measured haemoglobin levels in the brain tissue (not shown), and indeed reports show that at least 4 hours of MCA occlusion is required to induce haemorrhagic transformation with tPA(62). Importantly, co-treatment with tPA and anti-VCAM-CD39 did not cause a measurable increase in haemoglobin levels in brain lysates. This shows that low-dose targeted CD39 does not promote systemic bleeding even when co-injected with tPA. The synergistic interaction between tPA-thrombolytic therapy and anti-VCAM-CD39 treatment observed herein in the absence of detectable intracerebral bleeding has significant therapeutic implications for ischaemic stroke.

Though overexpression of CD39 is not associated with increased bleeding complications, CD39 transgenic mice do exhibit impaired platelet aggregation, prolonged bleeding times and resistance to systemic thromboembolism(25–27). We did not expect systemic bleeding at the dose of 0.5mg/kg of *anti-VCAM-CD39* used here as it did not increase tail bleeding time at doses ≤1mg/kg (**Supplementary Figure C**). Almost 20% of strokes are haemorrhagic at presentation(63). Further, stroke is 17% more prevalent in regional areas(64) and yet current therapies cannot be delivered in these areas as neuroimaging is a prerequisite that is usually available only during working hours or requires long distance travel for access in rural and regional areas, particularly in Australia(65). To demonstrate that *anti-VCAM-CD39* can be safely administered without the need for imaging to determine whether the stroke is ischaemic or haemorrhagic, we tested its effect in a haemorrhagic stroke model. The collagenase injection model of ICH is most widely used and best recapitulates blood vessel rupture leading to bleeding, re-bleeding, and consequent haematoma expansion(66). Despite the well-defined and potent antithrombotic activity of CD39(25–27), *anti-VCAM-CD39* did not worsen bleeding or neurological dysfunction within 24 hours post-ICH. There was a significant reduction in caspase-3 activity in the injured hemisphere; it remains to be determined if lesion volume was altered. It is remarkable that *anti-VCAM-CD39* is neuroprotective even in instances of haemorrhage, with no further bleeding risk. Further, tail bleeding times were not prolonged with administration of *anti-VCAM-CD39* at doses ≤1mg/kg (Supplementary Figure C), like other studies where targeted CD39 has been utilised(28).

Taken together, we have shown that *anti-VCAM-CD39* protects the brain post-MCAo, due to its ability to diminish the pro-inflammatory effects of eATP and likely due to adenosine-mediated restoration of BBB permeability and microvascular perfusion. More in-depth analysis of the protective mechanism of *anti-VCAM-CD39* still needs to be done to confirm and understand its action in the ischaemic stroke. A caveat noted by us is the absence of a control construct that had both a scrambled scFV VCAM component and an inactive CD39 component; we recommend that further preclinical studies include this additional fourth control. We also note the reduced sample sizes for the MRI data due to the high cost of accessing this service. Similarly, due to limited resources and the large number of experiments, it was unfeasible for us to use both male and female mice at this stage. Future experiments should include similar analyses in female mice.

Widespread global inflammation, BBB damage and microvascular thrombosis are not addressed by current therapies. There is an unmet clinical need for new therapeutics that can target the inflammatory and ischaemic penumbra, restore microcirculation, and can be administered to patients at delayed timepoints post ischaemia without the need for prior imaging, which is particularly relevant for people in regional and rural areas. Further, a therapeutic that could synergise with existing thrombolytic therapies would have the ability to transform stroke treatment. *Anti-VCAM-CD39* appears to be safe for administration as a single agent, as shown here. The added benefit seen in combination with tPA creates a strong case for further development of this agent for the management of ischaemic stroke.

## Methods

### Design of the *anti-VCAM-CD39* expression construct and protein production

Anti-VCAM-1 ScFv was generated in collaboration with Prof Claudia Gottstein and Prof AI Karlheinz Peter and was translationally fused to CD39. Refer to supplementary methods for further information.

### *In vivo* experiments

#### Mice

Experiments were performed in male C57BL/6 mice (age, 8 ± 0.2 weeks; weight, 28.2 ± 2 g) obtained from an in-house colony where they were placed on a 12h light-dark cycle with *ad libitum* access to food and water.

#### MCAo

Transient focal cerebral ischaemia (30 minutes) was induced using MCAo, as described previously(42). Briefly, MCAo was induced by occluding the middle cerebral artery with a silicone rubber-coated monofilament suture for an ischemic period of 30 minutes. The common carotid artery was allowed to perfuse for the duration of ischemia.

#### ICH

Mice were anaesthetised with 2,2 tribromoethanol (Avertin) and 0.15U (1 µl) of Collagenase VII (from Clostridium *histolyticum*; both from Sigma Aldrich Australia) was injected intracerebrally as described(42, 67) at coordinates MV: 1.75, AP: 0.4, DV: 3.8 from bregma.

#### Treatment Administration

We had previously determined that VCAM-1 was significantly upregulated 3 hours(42). We tested anti-VCAM-CD39 at various doses (not shown) and determined that the optimum therapeutic dose was 0.5mg/kg, which we administered intravenously 3 hours post-surgery. The corresponding dose of the controls (non-targeted CD39; equi-active-1.5 mg/kg; and anti-VCAM-inactive CD39-0.5 mg/kg) was similarly administered. Mice were randomly assigned to experimental groups and given a sample number, and all mouse groups were blinded during subsequent assessment. Animals were left to recover on the heat pad for at least 4 hours after procedure. Animals were separated and given supportive treatment and left on the heat rack overnight(42).

#### tPA Administration

Recombinant mouse t-PA (0.9 mg/kg; Molecular Innovations, MI, USA) or saline (equi-volume) was co-administered 3 hours after reperfusion as a bolus intravenous injection via the tail vein.

#### Euthanasia and tissue harvesting

At 24 hours post-surgery, mice were anaesthetized with Lethabarb (Pentobarbitone, 90mg/kg, Australia), and transcardially perfused with phosphate buffered saline (PBS) pH 7.4. Unless otherwise stated, a 6mm section of the infarcted brains was dissected and homogenised to 300mg wet weight of tissue per 1ml of Lysis Buffer (TBS +1% Triton X-100; Roche Australia).

#### Functional Assessment

Neurological assessment was done through Bederson scoring as described previously(42, 68) 20-24 hours post-stroke.

#### Magnetic resonance imaging

At 24 hours post-stroke, *in-vivo* MRI Imaging was performed using a 9.4 T/20 cm Bruker MRI with actively decoupled volume transmit and surface-receive coils as described previously(69). Refer to supplementary methods for further information.

#### Western Blot

75μg of total protein was subjected to SDS-PAGE. Caspase-1 p20 and Actin levels in tissue lysates were detected by rabbit polyclonal anti-Caspase 1 p20 antibody (Thermo Fisher, Australia) and rabbit HRP conjugated anti-beta-Actin (Cell signalling, Australia). 2μL of plasma was subjected to SDS-PAGE. vWF levels in plasma were detected by sheep monoclonal anti-vWF. Refer to supplementary methods for further information.

#### Determination of plasma ATP levels

Refer to supplementary methods for further information.

#### Measurement of plasma adenosine

The Adenosine Assay kit was used (Fluorometric) (#ab211094, Abcam, Australia) according to the manufacturer’s instructions. Refer to supplementary methods for further information.

#### Real-Time (RT)-PCR

Refer to supplementary methods for further information.

#### Detection of microglia via IBA1 immunofluorescence

Refer to supplementary methods for further information.

#### Blood brain barrier permeability assay

Albumin content in the brain was determined using the Mouse Albumin ELISA Quantitation Set (Bethyl Laboratories, USA) according to the manufacturer’s instructions and as described(42, 70). Albumin in the brain parenchyma is reflective of the extent of BBB damage as published(70). Refer to supplementary methods for further information.

#### Statistics

was performed using Prism 9 software (GraphPad, US). Normality tests were run to determine subsequent statistical test. Confirming normality, comparison of experimental datasets was performed by one-way ANOVA with Dunnett’s post-hoc correction or two-way ANOVA with Sidak’s or Dunnett’s post-hoc correction as stated. Non-normal datasets were compared with Kruskal-Wallis test with Dunn’s multiple comparisons test. Differences between two groups were assessed by two-tailed student t-tests (unpaired or unpaired with Welch’s correction for parametric data and Mann-Whitney test for non-parametric data). P< 0.05 was considered significant.

#### Study Approval

All experiments were approved by Alfred Research Alliance Ethics Committee (ethics applications E/1851/18/M and E/1937/19/M) in accordance with Australian code for the care and use of animals for scientific purposes 8^th^ Ed 2013 and in compliance with ARRIVE guidelines for reporting animal experiments.

#### Data availability

Values for all data points in graphs are reported in the Supporting Data Values file. Any additional data can be requested from corresponding author MS.

## Author contributions

NL: Conceptualization, Data curation, Formal analysis, Investigation, Methodology, Validation, Roles/Writing - original draft, Writing - review & editing.

IS, CS, IC, JC, AW: Data curation, Investigation, Methodology

DKW, RB: MRI-Data curation, Software, Investigation, Methodology

XW, KHP, SR, RLM-Conceptualization, Writing - review & editing.

HN: Conceptualization, Funding acquisition, Investigation, Methodology, Project administration, Resources, Supervision, Writing - review & editing.

MS: Conceptualization, Data curation, Formal analysis, Funding acquisition, Investigation, Methodology, Project administration, Supervision, Validation, Roles/Writing - original draft, Writing - review & editing.

## Acknowledgements

We acknowledge the Monash Histology Platform and the Monash Microimaging Platform, Monash University, for the provision of instrumentation, training and technical support. The authors acknowledge the facilities and scientific and technical assistance of the National Imaging Facility (NIF), a National Collaborative Research Infrastructure Strategy (NCRIS) capability at Monash Biomedical Imaging (MBI), a Technology Research Platform at Monash University.

This work was supported by NHMRC Project grant APP1141046 awarded to HN and Bethlehem Griffiths Research Foundation Grant (#1708) given to HN and MS.

